# Multimodal Single-Cell Characterization of the Human Granulocyte Lineage

**DOI:** 10.1101/2021.06.12.448210

**Authors:** Jingjing Qi, Darwin D’Souza, Travis Dawson, Daniel Geanon, Hiyab Stefanos, Robert Marvin, Laura Walker, Adeeb H. Rahman

**Author notes:** Corresponding author: Adeeb Rahman.

## Abstract

High throughput single cell transcriptomics (scRNA-seq) has been successfully applied to characterize immune cell heterogeneity across a diverse range of settings; however, analysis of human granulocytes remains a significant challenge due to their low gene expression transcript detection. Consequently, granulocytes are typically either absent or highly under-represented and inaccurately enumerated in most human scRNA-seq datasets. Here, we apply multi-modal CITE-seq profiling to characterize granulocytes in human whole blood and bone marrow, and we show that these populations can be accurately detected and analyzed using the antibody-based modality, and that their frequencies and phenotype align well with antibody-based characterization of the same samples using CyTOF. These analyses also clearly highlight extremely low gene transcript detection across the entire granulocyte lineage including the earliest neutrophil progenitor populations when using the 10X Genomics platform. By contrast, when performing parallel analyses of the same samples using the BD Rhapsody platform, we recovered a much higher proportion of granulocyte gene transcripts, enabling true multi-modal characterization of human granulocyte heterogeneity.

## Introduction

High-throughput single-cell transcriptomic (scRNA-seq) technologies are increasingly being applied to study immune cell heterogeneity across a wide range of diseases^1–4^. The 10X Genomics platform is currently one of the most widely utilized commercial platforms for high throughout scRNA-seq^5^. The platform utilizes microfluidics to encapsulate single cells and reverse transcription reagents with barcoded gel beads, enabling the generation of barcoded cDNA within each droplet. To ensure the pairing of single cells with beads, the system utilizes an excess of beads and introduces cells at a limiting dilution such that the vast majority of bead-containing droplets do not contain a cell. Thus, one of the first steps in performing scRNA-seq analysis is to filter out “empty” bead barcodes that were never paired with a cell to restrict analysis to only cell-associated barcodes. This is typically accomplished by thresholding on the gene UMI count of each barcode based on the assumption beads that were encapsulated with a cell will have high gene UMI counts while those that were not will have low UMI counts.

While the measurement of thousands of genes can resolve the complexity of phenotypic and functional heterogeneity, gene expression does not always serve as an adequate proxy for protein expression, and it can sometime be challenging to relate cell populations defined by unbiased transcriptomic clustering to those identified by historical antibody-based approaches. Recent methods, such as CITE-seq, have introduced the use of oligonucleotide labelled antibodies to enable simultaneous single cell detection of protein epitopes in parallel with unbiased transcriptome profiling^6, 7^. When performing multi-modal CITE-seq analyses, cell-associated barcodes can be defined by thresholding on either total gene UMIs or antibody UMIs. The latter approach is particularly relevant when utilizing antibodies against ubiquitous cell surface antigens for cell “hashing”^8^.

When analyzing high quality peripheral blood mononuclear cells (PBMCs) isolated using density centrifugation, we find that total gene and hashtag antibody UMIs are typically well correlated and that thresholding on either results in a very similar number of cell barcodes (Figure 1A). The number of cell barcodes aligned well with the estimated number of cells captured based on pre-encapsulation cell counts (Figure 1B). In these samples, the small number of cells with high hashtag UMIs but low gene expression UMIs represented low quality/viability cells. Conversely, the small number of cells with high gene UMIs but low hashtag UMIs were identified as red blood cells, which do not express high levels of the antigens detected by the hashing antibodies.

**Figure 1.**
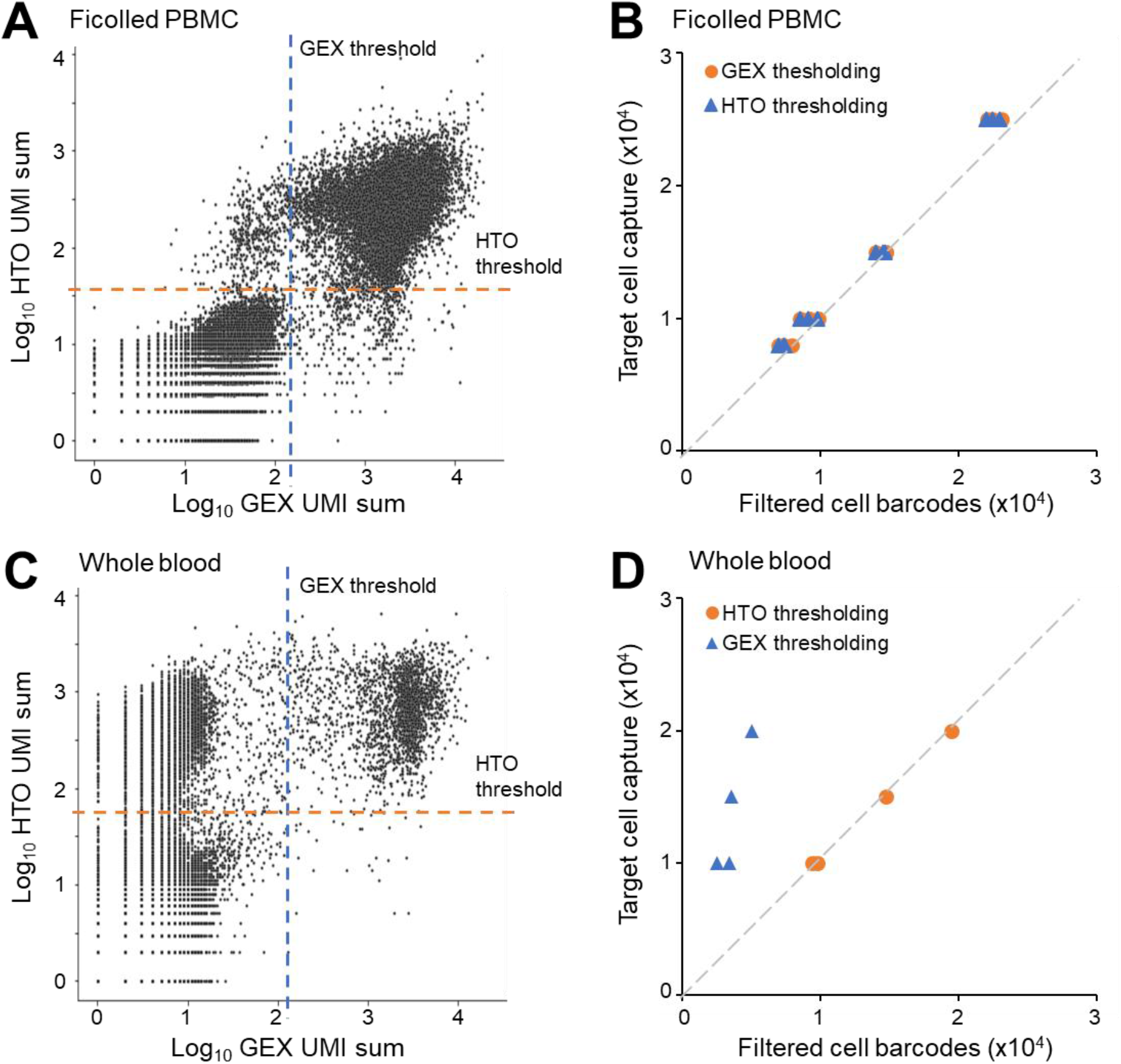
Discrepancies in cell barcode identification by gene and hashtag antibody UMIs in PBMCs and whole blood/bone marrow. (A) Representative biaxial plot of an unfiltered Cell Ranger matrix showing the correlation between total gene and HTO UMIs in a PBMC sample analyzed with the 10X Genomics 3’v2 kit. Dashed lines indicated thresholds applied to filter cell-associated barcodes from empty droplets. (B) Correlation between the number of filtered cell barcodes defined by either gene or HTO based thresholding and the number of cells targeted for capture based on pre-encapsulation cell counts. Each point represents a distinct PBMC sample. (C) Representative biaxial plot of an unfiltered Cell Ranger matrix showing the correlation between total gene and HTO UMIs in a RBC-depleted whole blood sample analyzed with the 10X Genomics 3’v2 kit. (D) Correlation between the number of filtered cell barcodes defined by either gene or HTO based thresholding and the number of cells targeted for capture based on pre-encapsulation cell counts. Each point represents a distinct whole blood or bone marrow sample.

However, when analyzing dissociated tissue cell suspensions and other sample types not processed using density centrifugation, such as RBC-depleted whole blood and bone marrow, we noted a high proportion of cells expressing high levels of hashtag antibody UMIs but low levels of gene UMIs (Figure 1C). These low-UMI cell barcodes are typically excluded by traditional gene UMI-based filtering as is typically performed in default Cell Ranger processing, resulting in a much lower recovery of cell barcodes than would be expected based on pre-loading cell counts (Figure 1D).

### CITE-seq analyses with the 10X Genomics platform allow for accurate enumeration and antibody-based characterization of human granulocytes but highlight low gene UMIs

We noted that the proportion of cells with divergent gene and hashtag antibody UMIs correlated directly with the expected proportion of granulocytes in these samples. Notably, while mouse granulocytes have been successfully characterized by scRNA-seq using the 10X Genomics platform^9–11^, granulocytes are largely absent or extremely under-represented in human data scRNA-seq datasets generated using the 10X Genomics platform^12, 13^. Despite their low gene expression, we reasoned that the robust detection of antibody-derived UMIs would allow for accurate identification and deeper phenotypic characterization of these cell populations using CITE-seq. To specifically focus on granulocytes, we analyzed fresh whole blood and bone marrow samples from healthy human donors, which were processed without gradient centrifugation to preserve all granulocyte populations. Following RBC and platelet depletion, the leukocytes were stained with a panel of 30 oligonucleotide-conjugated antibodies designed to identify and characterize most major immune cell types particularly granulocytes and granulocyte progenitors (Table 1). In addition to the phenotypic antibody panel, the samples were also split, stained with hashtag antibodies, and re-pooled prior to encapsulation to facilitate identification of cell-cell multiplets. The stained blood and bone marrow samples were analyzed using the 10X Genomics Chromium platform using the 3’v3.1 chemistry.

**Table 1.**
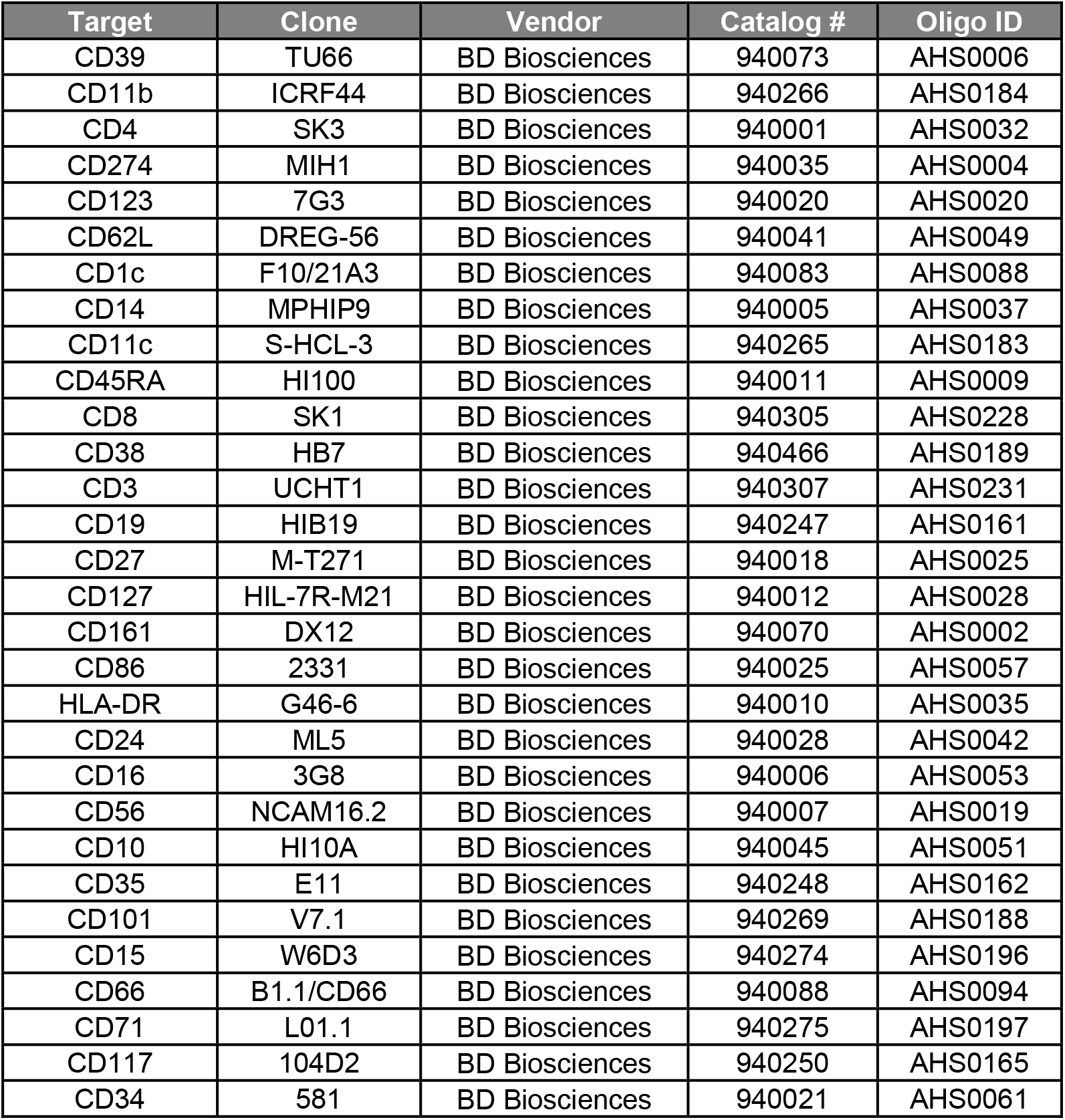
AbSeq antibodies.

As expected, the unfiltered Cell Ranger matrices derived from these samples contained a large proportion of cells with low gene UMIs so were instead processed by thresholding on antibody UMIs, resulting in a number of cell barcodes that closely matched the targeted cell recovery. The filtered cell barcodes were clustered based on antibody expression, and major immune cell types and progenitor populations were manually annotated based on canonical protein expression patterns (Figure 2A-C). Notably, our antibody panel enabled clear delineation of granulopoiesis including identification of granulocyte-monocyte progenitors (GMP), multipotent neutrophil progenitors, pre-neutrophils, immature neutrophils, mature neutrophils and aged neutrophils, which exhibited protein expression phenotypes consistent with other reports^14–17^.

**Figure 2.**
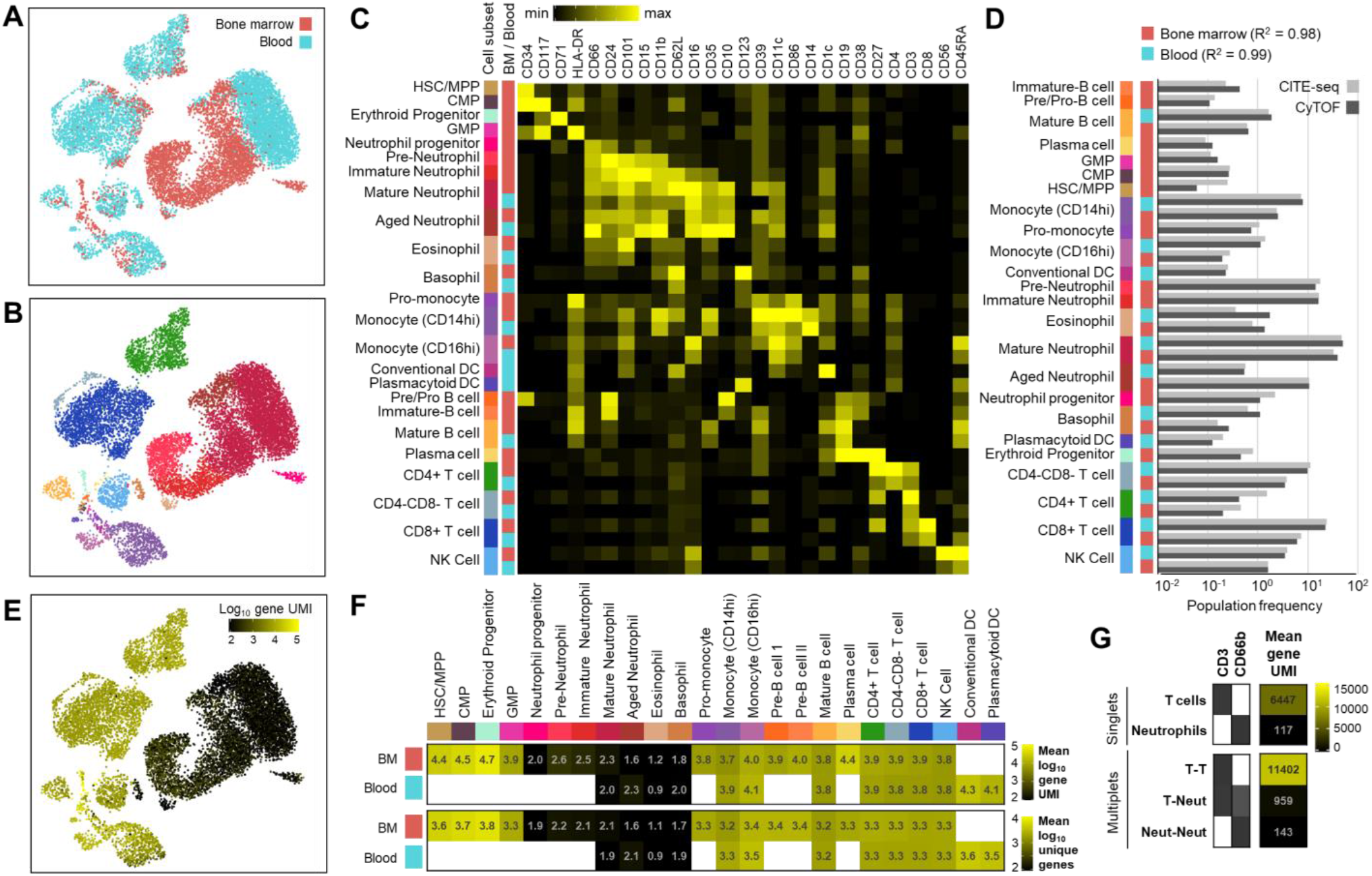
Multi-modal profiling of human peripheral blood and bone marrow with the 10X Genomics platform results in low gene detection in cells of the granulocyte lineage. Fresh whole blood and bone marrow samples from two healthy donors were depleted of RBCs, stained with a panel of 30 oligonucleotide-conjugated antibodies, and encapsulated using a 10X Genomics Chromium 3’v3.1 chip. Cell-associated barcodes were filtered from the resulting Cellranger matrix by thresholding on total antibody-transcript UMIs, and cell types were manually annotated based on protein expression of canonical cell surface markers. (A-C) UMAPs generated using antibody markers only and colored by sample source (A) or annotated cell type (B), with a corresponding hierarchically-clustered heatmap of scaled protein expression for each of the annotated subsets (C). (D) Parallel aliquots of the same RBC-depleted blood and bone marrow samples were analyzed by CyTOF using a matched antibody panel, and the relative frequencies of the paired cell subsets were compared across both platforms. (E) UMAP with cells colored by total log_10_ gene UMIs. (F) Mean log_10_ gene UMI (top) and unique genes (bottom) for each of the annotated cell subsets. (G) Cell singlets and known multiplets were defined using sample hashtags. Mean gene UMIs are shown for T cells singlets and multiplets (CD3+CD66b-), neutrophil singlets and multiplets (CD3-CD66b+) and in T cell-neutrophil multiplets (CD3+CD66b+).

To validate the cell annotations defined by our CITE-seq antibody-based annotations, parallel aliquots of the same RBC-depleted blood and bone marrow samples were stained with a panel of isotope-conjugated antibodies and analyzed by mass cytometry (CyTOF). Major immune cell types were similarly manually annotated using shared cell surface markers, and the relative proportion of cells was compared and found to be highly concordant between both platforms, supporting the fidelity of the CITE-seq antibody-based annotation (Figure 2D and Supplemental Figure 1). We next visualized the heterogeneity of transcriptional content across these annotated cell types (Figure 2E-F). Consistent with our hypothesis, we noted extremely low gene UMI and unique across all granulocyte populations, with mature neutrophils and eosinophils exhibiting gene UMIs and unique gene counts 10-100 fold lower than most other populations. Strikingly, while we observed high levels of gene UMIs in multipotent GMPs, the dramatic reduction of gene UMIs was observed at the earliest stage of granulocyte committed neutrophil progenitors. Low gene counts persisted throughout all stages of granulopoiesis with the lowest levels observed in aged neutrophils and eosinophils.

While these results suggested extremely low levels of mRNA in these granulocyte cell types, this did not seem consistent with prior bulk sequencing studies^18^. We instead hypothesized that the low gene detection in these cells may be due to high levels of nucleases that degraded mRNA transcripts during cell encapsulation and lysis. To test this hypothesis, we utilized cell hashing antibodies to identify neutrophil and T cell singlets, expressing CD66b and CD3 respectively, and only a single hashtag. We contrasted these cells with known cell-cell multiplets defined based on the expression of more than one hashtag antibody, which we further defined as T-T multiplets, neutrophil-neutrophil multiplets, or T-neutrophil multiplets based on co-expression of CD66b and CD3 (Figure 2G). A cell-cell doublet would generally be expected to contain the sum of UMIs from both contributing cells, and consistent with this, T-T cell multiplets typically showed a UMI count that was approximately 2-fold higher than T cell singlets. If neutrophils simply lacked mRNA content, a neutrophil-T cell multiplet would therefore be expected to contain a similar number of gene UMIs to a T cell alone. However, we instead found that the majority of T cell-neutrophil multiplets contained significantly lower UMI counts than T cells singlets, with the median UMI count of T-cell-neutrophil doublets being almost 10-fold lower than T singlets, thus supporting our hypothesis that nucleases released by neutrophils result in degradation of gene transcripts from a co-encapsulated T cell. Interestingly, our observation that early neutrophil progenitors also gene UMI detection suggests that the development of these cellular nucleases occurs prior to the formation of mature granules.

### The BD Rhapsody platform results in higher gene recovery from granulocytes and enables multimodal analysis of human granulopoiesis

Notably, the degradation of granulocyte transcripts occurs despite the presence of RNAase-inhibitors contained in the 10X encapsulation reagents. Furthermore, supplementing RBC-depleted whole blood samples with additional RNAase inhibitor cocktail prior to 10X encapsulation did not appreciably improve recovered UMIs from neutrophils or other cell types (data not shown). We therefore sought to determine whether this phenomenon was also observed in other scRNA-seq platforms. We used parallel aliquots of the same antibody-stained blood and bone marrow cells used for our 10X Genomics experiments and analyzed them using a whole transcriptome analysis kit on the BD Rhapsody platform, which utilizes a microwell array system for single cell encapsulation with capture beads^19^. The associated gene and antibody libraries were sequenced at a comparable depth to the 10X libraries and the resulting cell-barcodes were clustered and annotated based on antibody expression as with the earlier 10X analysis. When evaluating transcriptional heterogeneity across the similarly annotated cell subsets, we noted that while gene UMIs and unique gene counts were still somewhat lower in granulocytes, the counts were significantly higher than observed with the 10X platform (Figure 3A). A titration of sequencing read depth showed that while the 10X 3’v3.1 chemistry resulted in approximately 5-fold higher gene UMIs for a given average reads per cell for most immune cell subsets, such as T cells, neutrophil gene UMIs were approximately 10-fold higher on the BD Rhapsody platform (Figure 3B).

**Figure 3.**
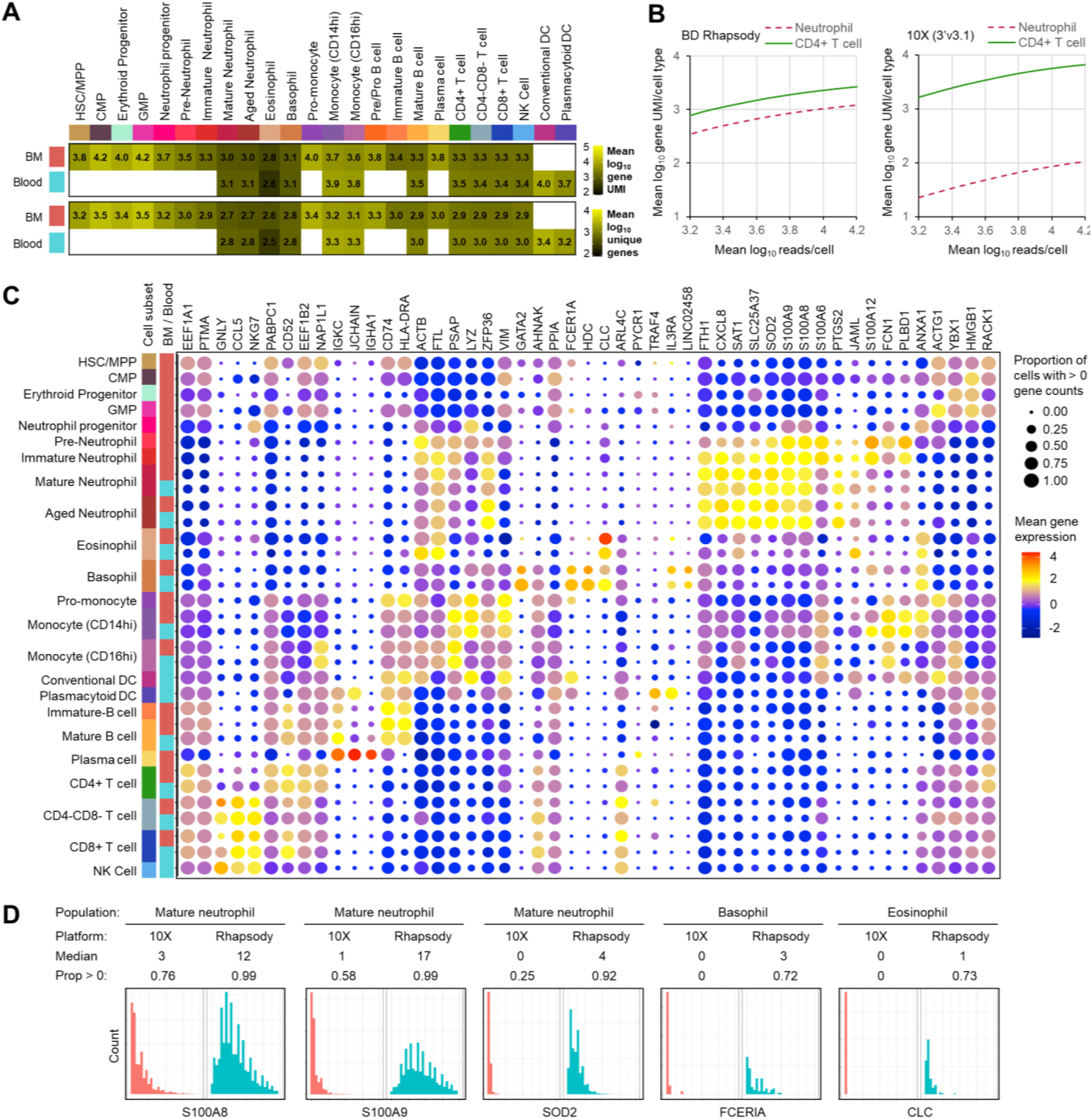
Multimodal profiling with the BD Rhapsody platform results in higher gene detection within the granulocyte lineage. Parallel aliquots of the same antibody-labeled blood and bone marrow samples shown in Figure 1 were encapsulated using the BD Rhapsody platform, and the same cell populations were similarly manually annotated using antibody expression alone. (A) Mean log_10_ gene UMI (top) and unique genes (bottom) for each of the annotated cell subsets. (B) Mean log_10_ gene UMI for blood neutrophils and CD4+ T cells at different read depths using the BD Rhapsody (left) and the 10X Genomics Chromium 3’v3.1 (right) platforms. (C) Mean expression and detection proportion of primary differential expressed genes for each of the annotated cell subsets. (D) Histograms comparing expression of highly expressed canonical genes in whole blood neutrophils, eosinophils and basophils using the BD Rhapsody and 10X Genomics platforms. The median expression and proportion of cells with non-zero gene counts are shown above each plot.

Given the higher gene recovery on the Rhapsody platform, we leveraged the multimodal data to integrate the data from both the blood and bone marrow and leveraged a recently described weighted nearest neighbor approach to analyze cells in both antibody and transcriptional space. This approach allowed us to clearly define key differentially expressed genes associated with distinct granulocyte lineages (Figure 3C), including high expression of several inflammatory S100 proteins and the chemokine CXCL18 in neutrophil subsets. Eosinophils and basophils expressed high levels of CLC, which encodes the Galectin-10 enzyme, and basophils additionally expressed FCERIA, GATA2 and HDC, which encodes histidine decarboxylase, an enzyme responsible for catalyzing the decarboxylation of histidine to form histamine. While single cell gene expression patterns were generally well conserved between the Rhapsody and 10X platforms, the lower gene capture resulted in lower mean expression levels and much higher levels of gene drop out even for canonical high abundance genes in the granulocyte populations in the 10X data (Figure 3D and supplementary Figure 2). Consequently, while gene expression based-integration of the datasets worked well for most immune populations, it was less effective for accurately resolving the heterogeneity amongst granulocyte populations (supplementary Figure 2).

We further leveraged the higher gene recovery on the Rhapsody platform to evaluate progressive gene expression changes associated with neutrophil maturation. We performed a gene-expression based clustering of all neutrophils in the bone marrow sample and found that the pseudotime trajectory estimated from the gene expression clustering broadly aligned with the antibody-based annotations (Figure 4A and B). Evaluating differential gene expression changes along the pseudotime trajectory revealed a clear signature associated with early neutrophil progenitors and a continuum of changes during maturation. The trajectory terminated with a subpopulation of neutrophils expressing CD83 together with high levels of the chemokines CCL3, CCL3L1, CCL4 and CCL4L1, suggesting a more inflammatory state. While the current study focuses only on samples from healthy donors, it is intriguing to consider whether this specific subpopulation may be expanded in the setting of infection^20^.

**Figure 4.**
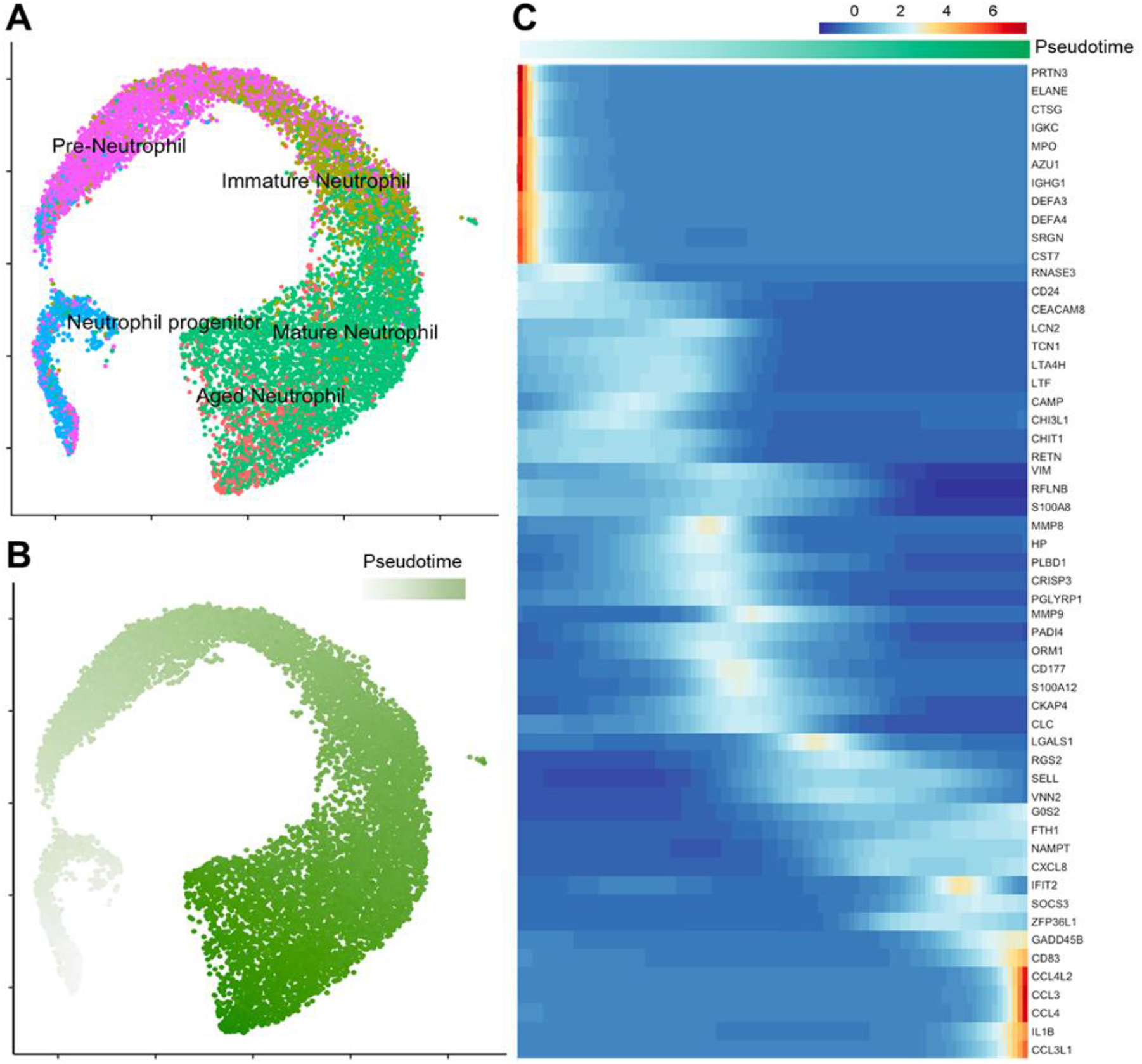
Trajectory-based differential gene expression analysis of human neutrophil development. (A-B) Gene expression UMAPs of bone marrow neutrophils analyzed using the BD Rhapsody platform colored based on antibody-based annotations (A) or estimated pseudotime based on gene expression clustering (B). (C) Estimated gene expression changes along the pseudotime trajectory.

Granulocytes play critical roles across many diseases ranging from allergy to systemic inflammatory responses to bacterial and viral sepsis, and scRNA-seq offers an opportunity to better characterize granulocyte heterogeneity and decipher the molecular mechanisms underlying these responses. However, our findings highlight the major limitations associated with studying human granulocytes using single cell transcriptional approaches on the 10X Genomics platform. While the incorporation of oligonucleotide-conjugated antibodies does allow for identification and characterization of granulocytes populations on the 10X Genomics platform based on their cell surface protein expression, our results the suggest that the BD Rhapsody platform is better suited to performing true single cell multi-modal analyses of granulocyte populations.

## Methods

### Samples

Fresh whole blood was collected from a healthy donor on the day of the experiment under an institutionally-approved IRB protocol. Fresh whole bone marrow was purchased from AllCells Inc. and delivered on the day of the experiment. The bone marrow was in a volume of 3mL and the sample was limited to 2 units/donor. The blood and bone marrow samples were depleted of red blood cells using StemCell Technologies EasySep™ RBC Depletion Reagent protocol. We found that this magnetic depletion method was superior to hypotonic lysis methods in preserving granulocyte quality. Platelets were subsequently depleted from the sample by a 5 minute centrifugation at 200 rcf. The remaining cells were resuspended in staining buffer (1x PBS + 0.2% BSA), counted and assessed for viability with acridine orange and propidium iodide staining using the Nexcelom Cellometer Auto2000 (Nexcelom Bioscience, Lawrence, MA, USA).

### Antibodies

Oligonucleotide-conjugated AbSeq antibodies for cell characterization were purchased from BD Biosciences (Table 1). Cell hashing antibodies targeting human Beta-2-microglobulin (clone 2M2) and CD298 (clone REA217) were purchased from Biolegend and Miltenyi Biosciences, respectively, and conjugated in house using Thunder-Link Plus Oligo Antibody conjugation kits (Expedeon) in accordance with the manufacturer’s instructions using an antibody:oligo ratio of 1:5. After conjugation, free oligo was depleted using 10 washes in a 50kDa Amicon filter. Isotope-conjugated antibodies for CyTOF analysis (Table 2) were either purchased pre-conjugated from Fluidigm or conjugated in-house using Fluidigm X8 polymer conjugation kits in accordance with the manufacturer’s protocols.

**Table 2.**
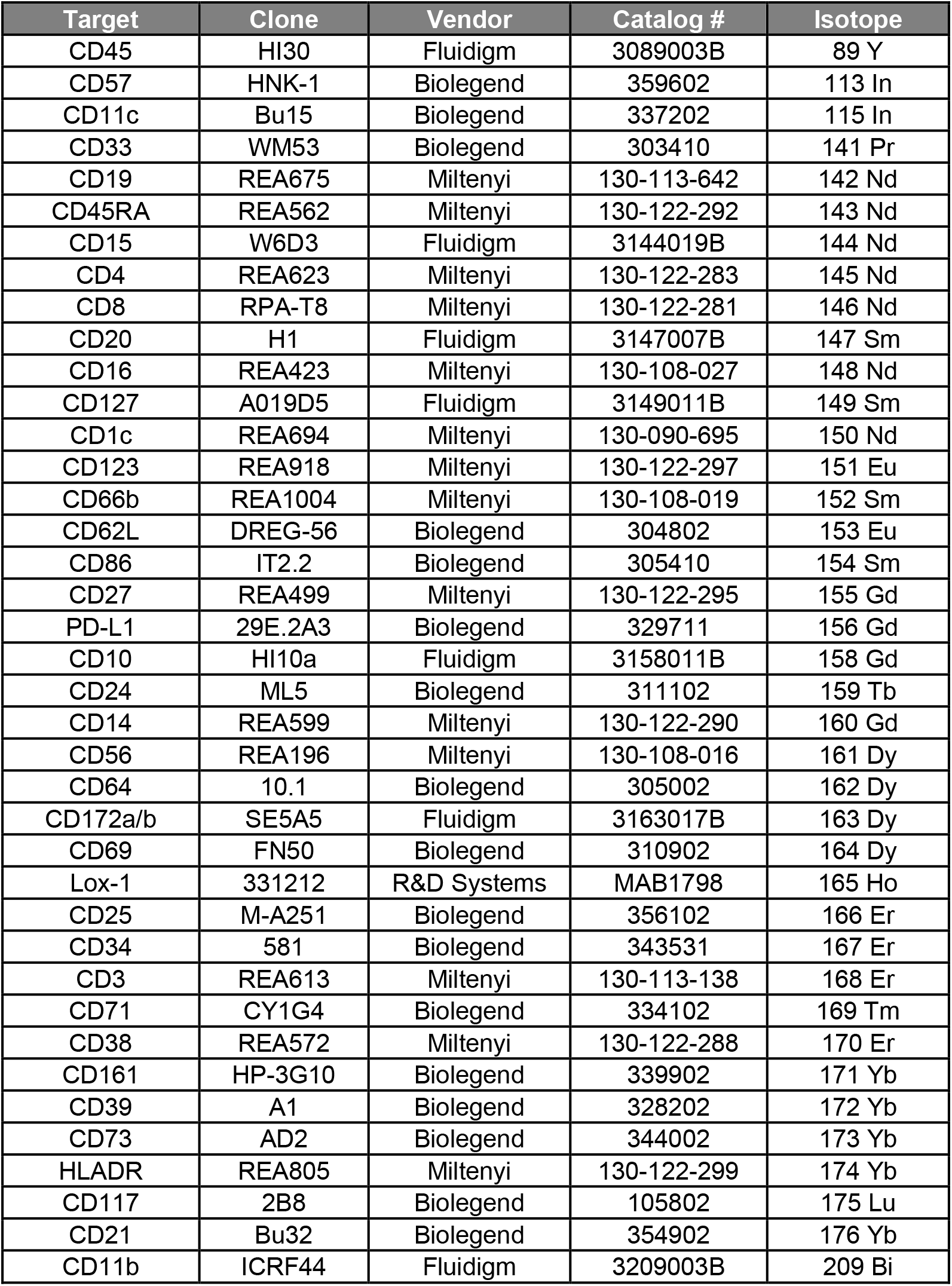
CyTOF antibodies.

### AbSeq antibody staining

Bone marrow and whole blood leukocytes were treated with human TrueStain FcX (Biolegend) to reduce non-specific antibody staining. The samples were first hashed by dividing into 10 aliquots of 250,000 cells each and stained with 10 hashtag antibodies at room temperature for 20 minutes. After washing three times by centrifugation at 350 rcf, the hashed samples were pooled and counted. 1 million hashed cells from the bone marrow and whole blood pools were then stained with a panel of 30 AbSeq antibodies at a dilution of 1:100 in a volume of 100uL. Residual antibodies were again washed 3 times to remove residual antibodies and counted using a Nexcelom Cellometer Auto2000 prior to encapsulation.

### 10X Processing

The hashed and stained bone marrow and whole blood pools were each loaded on one lane of the 10X Genomics Chromium Next GEM Single Cell 3’ v3.1 assay with a targeted cell recovery of 10,000 cells. Gene expression libraries were made as per the 10x Genomics demonstrated protocol. During cDNA amplification, primers designed to amplify the hashtag (GTGACTGGAGTTCAGACGTGTGC*T*C) and AbSeq (AATGATACGGCGACCACCGAGATCTACACTCTTTCCCTACACGACGC*T*C) primers were included in the PCR reaction to amplify the hashtag and AbSeq oligonucleotides respectively, and these amplicons were isolated from the cDNA amplicons via SPRISelect size selection. Hashtag oligo (HTO) libraries were prepared according to the New York Genome Center Technology Innovation Lab specifications “hashing” protocol. AbSeq libraries were prepared according to the New York Genome Center Technology Innovation Lab specifications “ADT” protocol. All libraries were quantified on the Agilent 2100 hsDNA BioAnalyzer and KAPA library quantification kit (Roche Cat# 07960140001). The resulting gene expression libraries were sequenced on an Illumina sequencing platform with a 28 × 8 × 60 bp configuration at a targeted depth of 25000 reads per cell, AbSeq libraries at a depth of 5000 reads per cell and HTO libraries at a depth of 1000 reads per cells.

### BD Rhapsody processing

Aliquots of the same hashed and stained bone marrow and whole blood samples used for the 10X experiment described above were each loaded in parallel onto a BD Rhapsody cartridge with a target capture of 20,000 cells each. Since cell capture and cDNA synthesis was performed using the BD Rhapsody Express Single-Cell Analysis System, and antibody and gene expression libraries were prepared using the AbSeq and Whole Transcriptome Analysis Amplification Kits, respectively. We were unable to successfully generate libraries for our in-house hashing antibodies using the Rhapsody platform. The resulting gene expression and AbSeq libraries were sequenced on an Illumina sequencing platform with 75 × 8 × 75 bp configuration a at a target depth of 25000 reads per cell and 5000 reads per cell, respectively to match the conditions used for the 10X analysis.

### CyTOF antibody staining

Aliquots of the same RBC-depleted bone marrow and whole blood leukocytes used for AbSeq staining were stained in parallel for CyTOF analysis. The cells were live-cell barcoded with cadmium-conjugated antibodies targeting Beta-2-microglobulin and CD298^21^, thus matching the same approach used for cell hashing. During the 30-minute live cell barcoding incubation, the cells were supplemented with Human TruStain FcX (Biolegend) to reduce non-specific antibody staining and 5uM Rhodium-103 (Fluidigm) for initial viability staining. After live-cell barcoding, cells were washed twice in flow buffer to remove residual barcode and pooled into one cell pool. After pooling, the cells were spun down and resuspended in the CyTOF antibody cocktail for 30 minutes at room temperature. Cells were then washed twice to remove residual CyTOF antibody and then subjected to Cell-ID Cisplatin (Fluidigm, San Francisco, CA, USA) post-stain viability staining following standard manufacturer protocols. Finally, the cell pool was fixed with 2.4% paraformaldehyde for 30 minutes at room temperature. 125nM Iridium-193 DNA intercalation and osmium tetroxide cell membrane labeling were performed simultaneously with cell fixation. Cells were then washed twice with staining buffer to remove excess iridium and osmium. Samples were stored at −80°C in FBS + 10% DMSO to preserve sample staining integrity^22^.

### CyTOF data acquisition / processing

Immediately prior to data acquisition, samples were thawed and room temperature and washed with Cell Staining Buffer and Cell Acquisition Solution (Fluidigm, San Francisco, CA, USA) and resuspended at a concentration of 1 million cells per ml in Cell Acquisition Solution containing a 1:20 dilution of EQ Normalization beads (Fluidigm, San Francisco, CA, USA). The samples were then acquired on a Helios Mass Cytometer equipped with a wide-bore sample injector at an event rate of <400 events per second. After acquisition, repeat acquisitions of the same sample were concatenated and normalized using the Fluidigm software. From here, CyTOF data processing and EQ bead based normalization was conducted with the provided Fluidigm software. Routine sample demultiplexing was conducted with the Zunder Lab debarcoder^23^. The demultiplexed files were then uploaded to Cytobank for subsequent sample clean-up. Specifically, EQ beads (140Ce+) and bead-cell doublets (140Ce+Ir193+) were excluded from the data, as well as Gaussian doublets (Residual high / Offset high) and cross-sample barcode doublets (barcode_separation_dist low / mahalanobis_distance high). Viable immune cells were identified as CD45+Ir+ and populations were annotated as described below.

### Data processing and analysis

BCL files for both 10X and Rhapsody samples were base-called and demultiplexed using cellranger mkfastq v3.1.0. For 10X samples, the alignment of gene expression libraries to human reference GRCh38 as well as the counting of unique features of the AbSeq and HTO libraries were performed using cellranger count v3.1.0. Similarly, Rhapsody samples were processed using BD Rhapsody™ WTA Analysis Pipeline, v1.9.1.

Putative cells in 10X samples were determined by gating the droplets gene umi sum and hashtag umi sum and choosing a threshold where the density was the highest. Cell-cell multiplets were identified based on co-expression of multiple hashing antibodies. This method could not be applied to the Rhapsody samples due to insufficient counts for the hashtags returned from the pipeline. Hence, putative cells were determined using Rhapsody’s second derivative analysis algorithm on the gene expression. Notably, in both cases we applied an inclusive thresholding strategy to include cell barcodes associated with either gene or antibody UMIs to ensure inclusion of granulocyte subsets. Multiplets and cells with more than 15% mitochondrial genes were excluded from downstream analyses unless specifically noted.

The 10X antibody staining data were normalized and denoised using DSB and visualized using Clustergrammer2 (https://github.com/ismms-himc/clustergrammer2)^24^. Phenotypically homogenous cell populations were identified by hierarchical clustering using the Clustergrammer2 interactive dendrogram and manually annotated based on canonical protein expression patterns. A consistent cell annotation strategy was applied across the 10X, Rhapsody and CyTOF antibody datasets to enable cross-platform comparisons.

Gene expression data were analyzed using the Seurat 4.0 R packages^25^. To compare the relative gene expression matrices generated from the different technology platforms (CITE-seq and Rhapsody) and tissue sources (blood and bone Marrow), we harmonized the gene expression datasets. Each individual dataset was log normalized and scaled followed by PCA. 3,000 most variable genes were used to identify the pairwise anchors of the datasets. We used the CCA method which has been shown to be able to effectively integrate datasets with shared biological markers and conserved gene correlation patterns even with presence of extensive technical and/or biological differences^26^. After data integration, primary differentially expressed genes between the annotated populations were determined using the presto R package^27^. For a given cell type, genes were selected by their capability to distinguish from other cell types (AUC > 0.7) and their mean expression (Top 5).

We explored the pseudotime trajectories in different stages of bone marrow neutrophil lineage. PCA was first performed on the gene expression of all neutrophil populations using the top 5000 variable geens, followed by generating the two dimensional projection in UMAP space using the first 50 principle components. 4 clusters were identified using Seurat’s graph-based clustering method. The Slingshot R package^28^ was used to estimate the pseudotime trajectories/potential lineages based on the UMAP embedding and cluster labels. Top 10 genes up or down regulated in adjacent cluster contrast were identified by (FDR < 0.05 and sorted by logFC). The R package tradeSeq^29^ was used to estimate gene expression along the trajectories.

### Data Availability

All data associated with this publication have been made publicly available. CyTOF FCS files are available on Flow Repository (https://flowrepository.org/id/FR-FCM-Z3T8).

For Multi-modal 10X and Rhapsody data, raw FASTQ files have been deposited into NCBI with BioProject ID: PRJNA734283. Raw Gene Expression and Antibody Stained matrices can be found at the following link: https://himc-project-data.s3.amazonaws.com/ADRA_RnD/sent_out/Neutrophil_BoneMarrow_WholeBlood_10X_Rhapsody.tar.gz

The integrated 10X and Rhapsody data are accessible at Galaxy^30^ via url: https://usegalaxy.eu/u/qij01himc/h/rhapbmneut. Interactive Environments can be created from the shared dataset to visualize the Integrated Multi-modal UMAP.

---

**Supplementary Figure 1.**
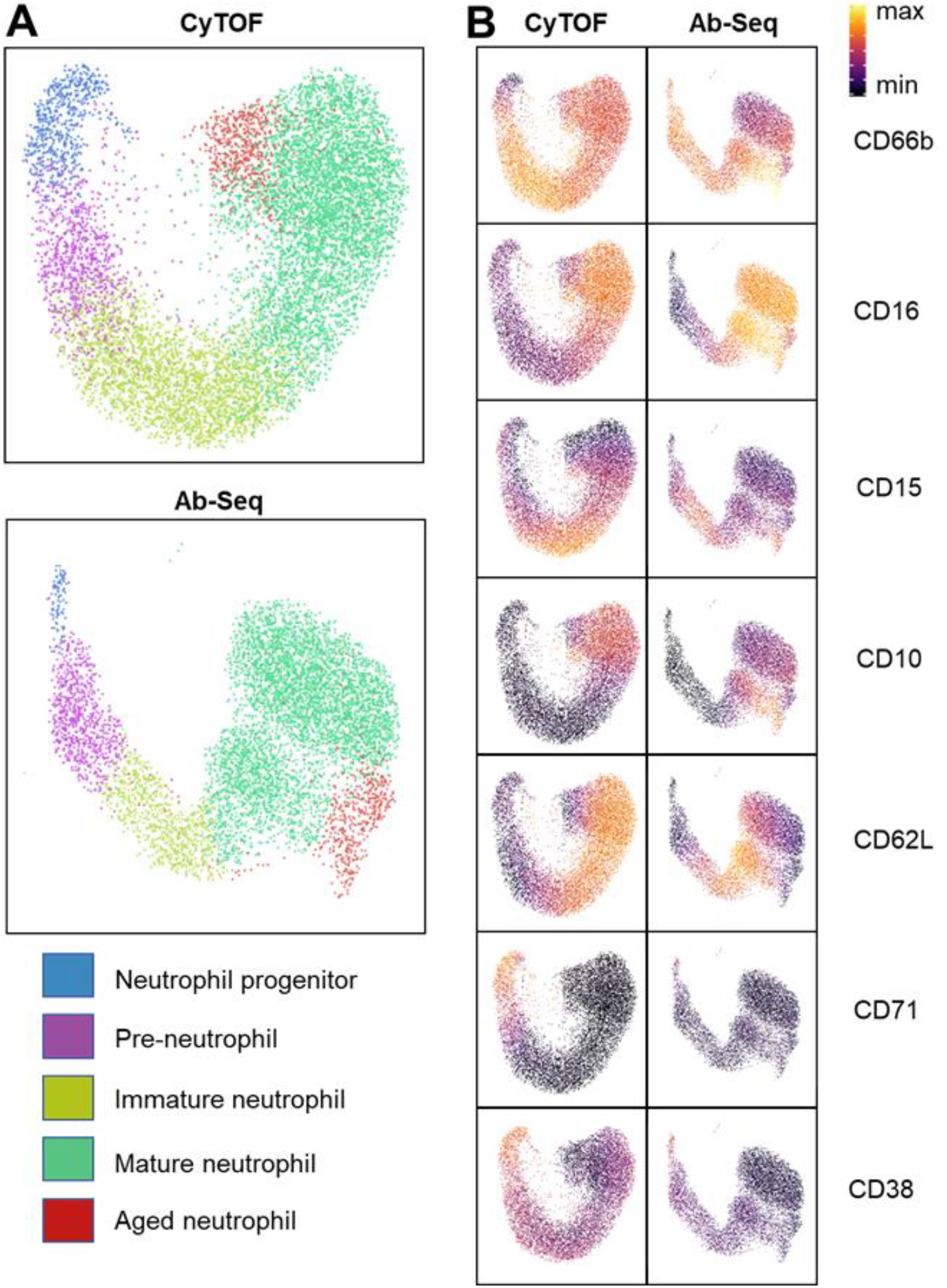
Representative CyTOF and AbSeq antibody staining on bone marrow neutrophils. UMAPs of bone marrow neutrophils analyzed by CyTOF (left) and AbSeq antibody staining (right). colored by (A) annotated population and (B) protein expression reveal relative concordance between the two immune profiling technologies.

**Supplementary Figure 2.**
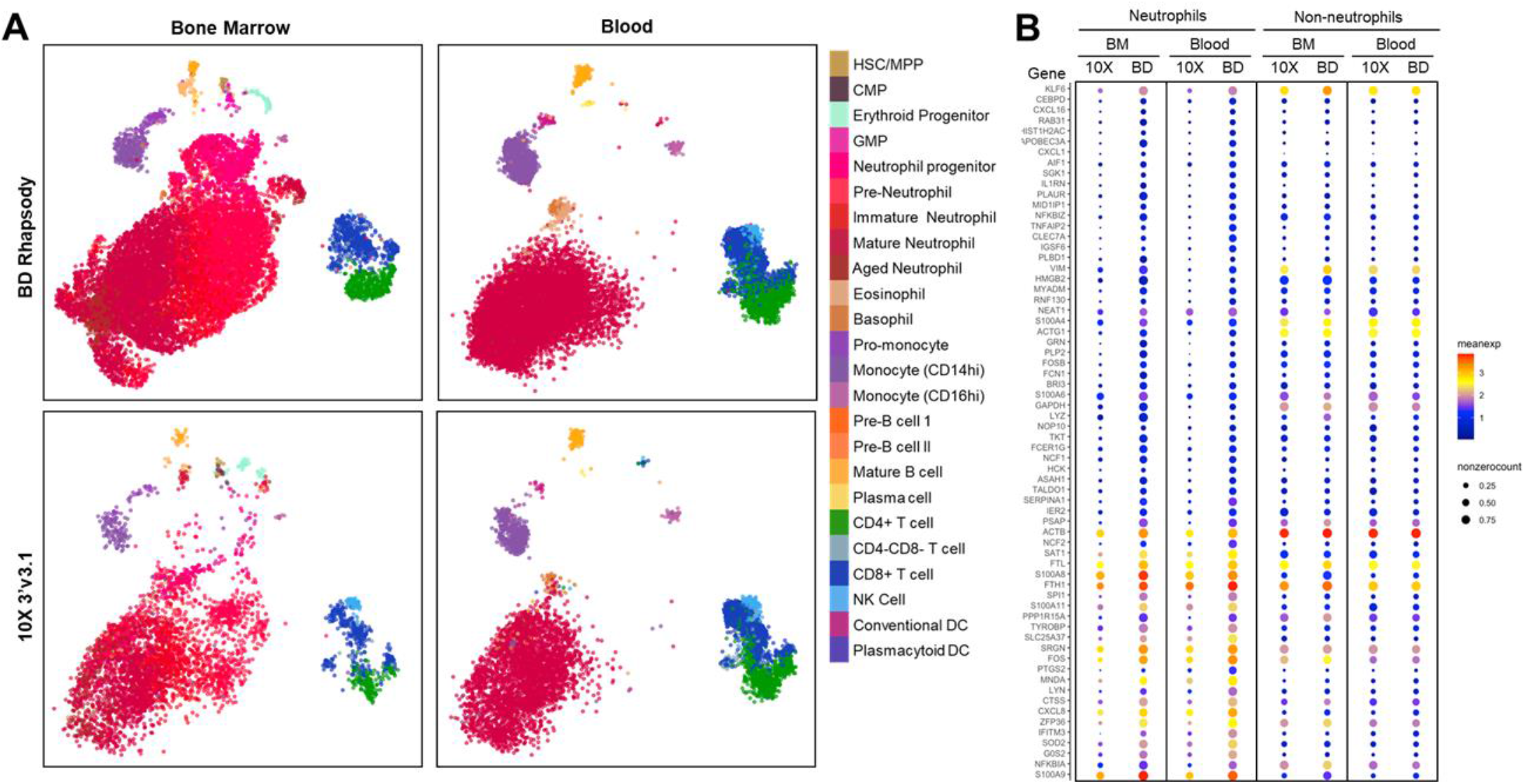
Gene expression integrated across platforms and sample types. (A) Post-integration UMAPs clustered by gene expression and colored by antibody-based cell type annotations. (B) Mean and non-zero gene counts for neutrophils and non-neutrophils for each of the platforms and sample types.

